# Ventrointermediate thalamic stimulation improves motor learning in humans

**DOI:** 10.1101/2023.09.19.558378

**Authors:** Angela Voegtle, Laila Terzic, Amr Farahat, Nanna Hartong, Imke Galazky, Hermann Hinrichs, Slawomir J. Nasuto, Adriano de Oliveira Andrade, Robert T. Knight, Richard B. Ivry, Jürgen Voges, Matthias Deliano, Lars Buentjen, Catherine M. Sweeney-Reed

## Abstract

Ventrointermediate thalamic stimulation (VIM-DBS) modulates oscillatory activity in a cortical network including primary motor cortex, premotor cortex, and parietal cortex. Here we show that, beyond the beneficial effects of VIM-DBS on motor execution, this form of invasive stimulation facilitates production of sequential finger movements that follow a fixed sequence but not when the sequence is selected at random. These results highlight the role of thalamo–cortical activity in motor learning.

## Main

Motor learning has been associated with a network that includes the cerebellum, primary motor cortex (M1), basal ganglia, ventrointermediate nucleus of the thalamus (VIM), and parietal–frontal cortex^1–5^. This network is disrupted in patients with essential tremor (ET)^6,7^, and deep brain stimulation of the VIM (VIM-DBS) has been shown not only to alleviate tremor, but also can improve motor learning^4,8^. To date, the neural mechanisms associated with this improvement are not well understood. Here we investigate the impact of VIM-DBS on oscillatory activity in cortical nodes of the motor learning network during sequence learning using the serial reaction time task (SRTT)^9^. Modulation of oscillatory brain activity has been observed in the alpha^5,10,11^ and beta bands^12^ during motor sequence learning. For example, patients with Parkinson’s disease have enhanced and prolonged beta band suppression that correlates with impairments in motor sequence learning^12^. ET patients exhibit increased suppression in the alpha/beta band during movement, similar to Parkinson’s disease patients^6^. We hypothesized that oscillatory power is modulated by VIM-DBS in patients with ET, with reduction of pathological electrophysiological brain activity in the alpha/beta frequency. Given that sequence learning is modulated by VIM-DBS^4^, we predicted greater cortical power modulation while patients performed a learned sequence of finger movements compared to when they produced a series of random finger movements.

We examined reaction times and scalp EEG oscillatory activity during SRTT performance, comparing conditions in which the VIM-DBS was ON or OFF (Fig. 1). The task required the participants to make a series of finger responses based on the position of a stimulus which either followed a fixed, 12-element sequence or was selected in a pseudorandom manner. We used cluster-based permutation tests^13^ to examine oscillatory spectral power across all trials (learned and random), comparing DBS-ON with DBS-OFF. We then performed separate analyses for learned and random trials, contrasting the ON-OFF conditions at the end of training and over the course of learning. We focused on stimulus-locked activity, using artifact removal methods^14^ that enabled analysis of EEG data recorded during DBS.

**Fig. 1.**
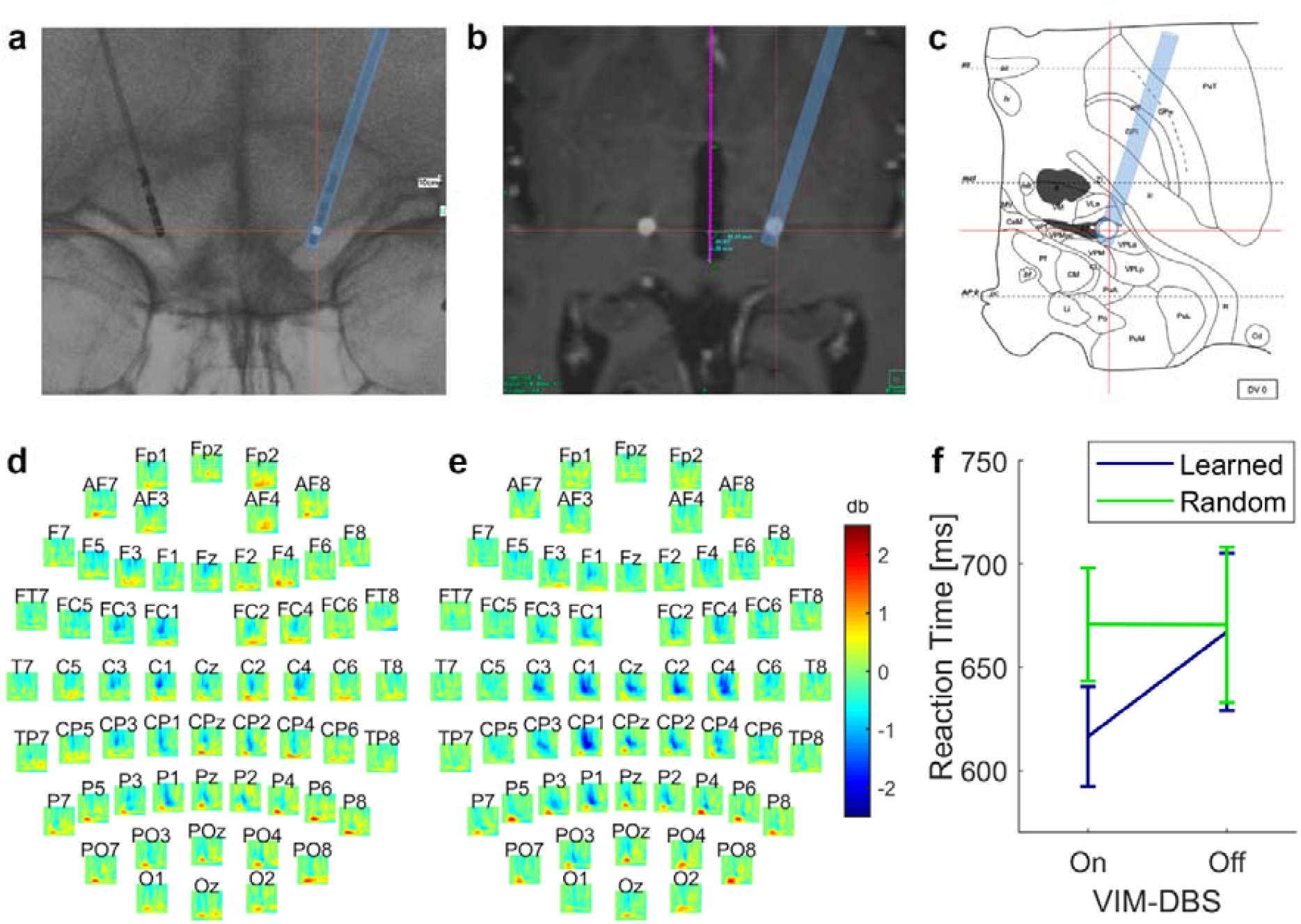
Illustration of stereotactic targeting of VIM. **a**. Stereotactic x-ray depicting intraoperative electrode location. Red crosshair: contact in the VIM (Vlpv-Morel^23^). **b**. Postoperative CT co-registered to preoperative MRI to establish electrode placement. White circle: CT-artifact of electrode contact in the VIM. Purple line: trajectory through the intracommissural line (IL). Red crosshair: electrode position in co-registered intraoperative x-ray. **c**. Stereotactic atlas of the thalamus^24^ at IL-level. Red crosshair: represents electrode position in relation to the IL. **d**. Grand-average during DBS-ON. **e**. Grand-average during DBS-OFF. **f**. Interaction for reaction times between *Stimulation Mode* (DBS-ON, DBS-OFF) and *Sequence type* (Learned, Random), showing mean and standard error. **d, e**: x-axis = Time: -200 to 1200 ms; y-axis = Frequency: 2-30 Hz.

In terms of behavior, there was an interaction between *Stimulation Mode* (DBS-ON, DBS-OFF) and *Sequence Type* (Learned, Random; F(1,11) = 8.75, p = 0.013, η^2^ = 0.44) Reaction times were faster during learned sequences compared to random sequences with DBS-ON (Learned: M = 616, 95% CI [563, 670]; Random: M = 671, 95% CI [611, 731]) compared to DBS-OFF (Learned: M = 667, 95% CI [583 751]; Random: M = 670, 95% CI [587, 754]) (T(11) = 2.96, p = 0.013, Cohen’s d = 0.85) (Fig. 1f).

Turning to the physiological data, a widespread, bilateral reduction in alpha/beta power suppression was seen when VIM-DBS was on compared with off (cluster-t = 9712, p_pos_ = 0.004, SD = 0.003, Cohen’s d = 2.156) (Fig. 1d-e). At the end of training when the stimuli followed a fixed sequence, power differences between DBS-ON and DBS-OFF encompassing the alpha/beta frequency bands were observed (Fig. 2). These effects were found over the central and ipsilateral motor area, including M1 and premotor cortex (PMC). During DBS-ON, alpha/beta power was less suppressed than during DBS-OFF in a cluster spanning a time window of ~320 ms, starting ~580 ms after stimulus onset (cluster-t = 5660, p_pos_ = 0.008, SD= 0.004, Cohen’s d = 1.779). No difference was detected between DBS-ON and DBS-OFF during random trials (cluster-t = 1407, p_pos_ = 0.325, SD = 0.021; cluster-t =-88, p_neg_ = 0.994, SD = 0.004). Post hoc evaluation confirmed an interaction between *Stimulation Mode* (DBS-ON, DBS-OFF) and *Sequence Type* (Learned, Random) (F(1,11) = 9.84, p = 0.009, η^2^ = 0.47). The power difference during learned sequences compared to random sequences was greater with DBS-ON (Learned: M = 0.22, 95% CI [-0.18, 0.62]; Random: M = -0.38, 95% CI [-0.74,-0.02]) compared to DBS-OFF (Learned: M = -0.95, 95% CI [-1.39, -0.51]; Random: M = -0.69, 95% CI [-1.19, -0.20]) (T(11) = 3.14, p = 0.009, Cohen’s d = 0.91) (Fig. 2c). Note a reduction in alpha suppression is distinct from an increase in alpha power; the former is less suppression, a recognized electrophysiological feature in motor cortex in patients with tremor.

**Fig. 2:**
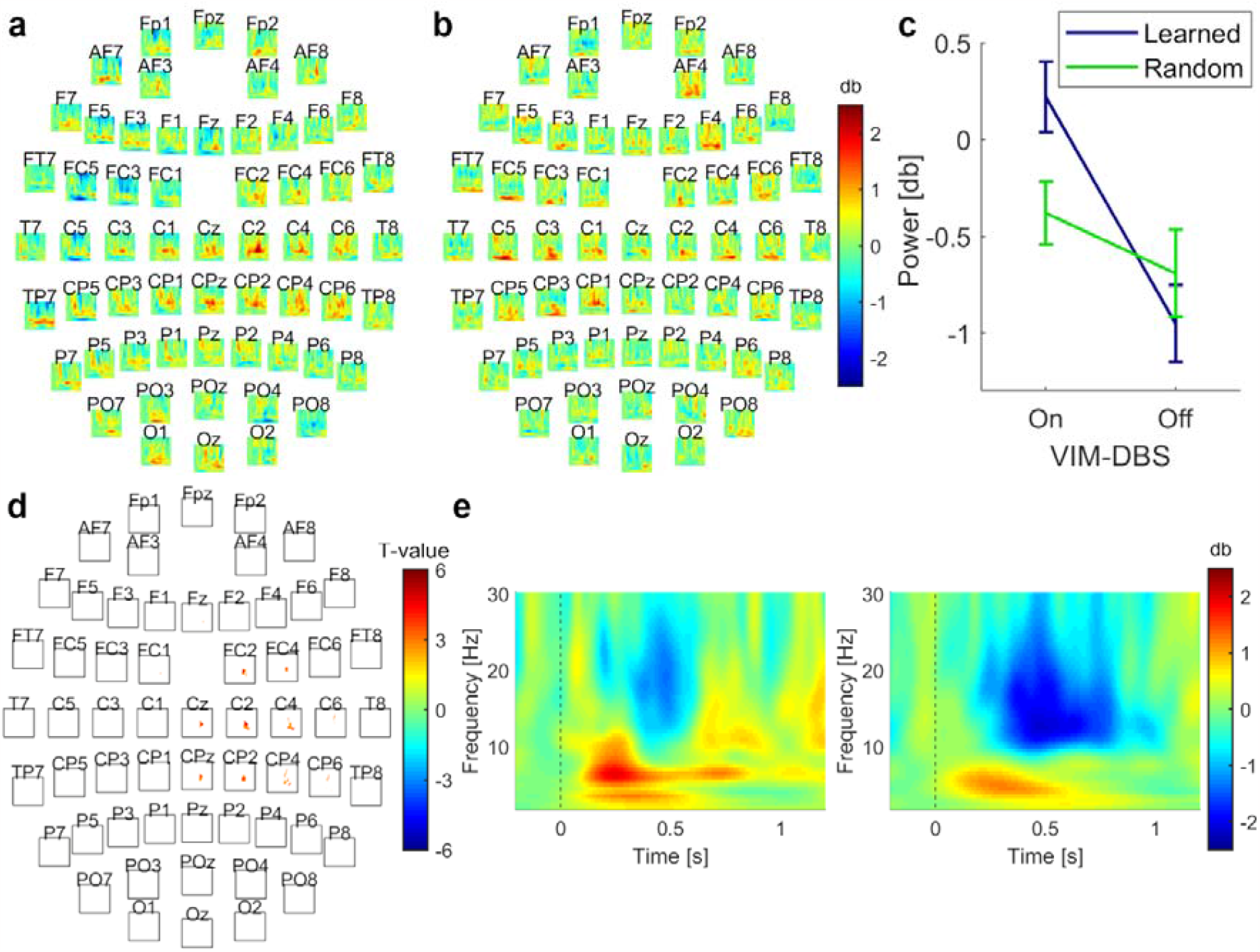
Oscillatory power during production of learned and random motor sequences (data from Block 4). **a** Grand-average difference between DBS-ON and DBS-OFF for learned sequences. **b** Grand-average difference between DBS-ON and DBS-OFF for random trials. **c** Interaction for power values between *Stimulation Mode* and *Sequence Type*, showing mean and standard error. **d** Location of the observed cluster, representing significant difference between DBS-ON and DBS-OFF during learned sequences. The cluster included electrodes spanning the ipsilateral sensorimotor cortex. The channel contributing most to the cluster was C2. **e** Left panel: channel C2 during DBS-ON; right panel: channel C2 during DBS-OFF. **a, b, d**: x-axis = Time: -200 to 1200 ms; y-axis = Frequency: 2-30 Hz.

Multiple linear regression was used to test whether the location of VIM-DBS stimulation predicted mean power over time, frequency, and EEG channel within the cluster, taking into account stimulation intensity and tremor severity. The overall regression was significant (R^2^ = 0.886, F(5,65) = 9.319, p = 0.009). Mean power was predicted by the AC–PC-x and AC–PC-y coordinates of the VIM stimulation contact, the total electrical energy delivered to the VIM, and tremor score, but not by the AC–PC-z coordinate (Supplementary Table 1). The power was greater when the VIM electrode was more lateral and anterior.

We also observed a difference in alpha power between DBS-ON and DBS-OFF late compared with early in learning (cluster-t = 7237, p_pos_ = 0.012, SD = 0.005, Cohen’s d = 2.485). The cluster was present over a central–posterior–occipital area contralateral to movement, starting ~500 ms after stimulus onset and persisting until 800 ms after stimulus onset; at some channel locations, the differences extended up to 1100 ms after stimulus onset (Fig. 3). Again, no difference was detected between the ON and OFF conditions during random trials (cluster-t = 2284, p_pos_ = 0.140, SD = 0.016; cluster-t = -241, p_neg_ = 0.884, SD = 0.014). Post hoc evaluation confirmed an interaction between *Stimulation Mode* (DBS-ON, DBS-OFF) and *Sequence Type* (Learned, Random; F(1,11) = 10.06, p = 0.009, η^2^ = 0.48). The increase in power during learned sequences compared to random sequences was greater with DBS-ON (Learned: M = 0.86, 95% CI [0.49, 1.23]; Random: M = -0.011, 95% CI [-0.51, 0.49])compared to DBS-OFF (Learned: M = -0.48, 95% CI [-0.80, -0.171]; Random: M = -0.26,95% CI [-0.59, 0.066]) (T(11) = 3.17, p = 0.009, Cohen’s d = 0.92) (Fig. 3c).

**Fig. 3:**
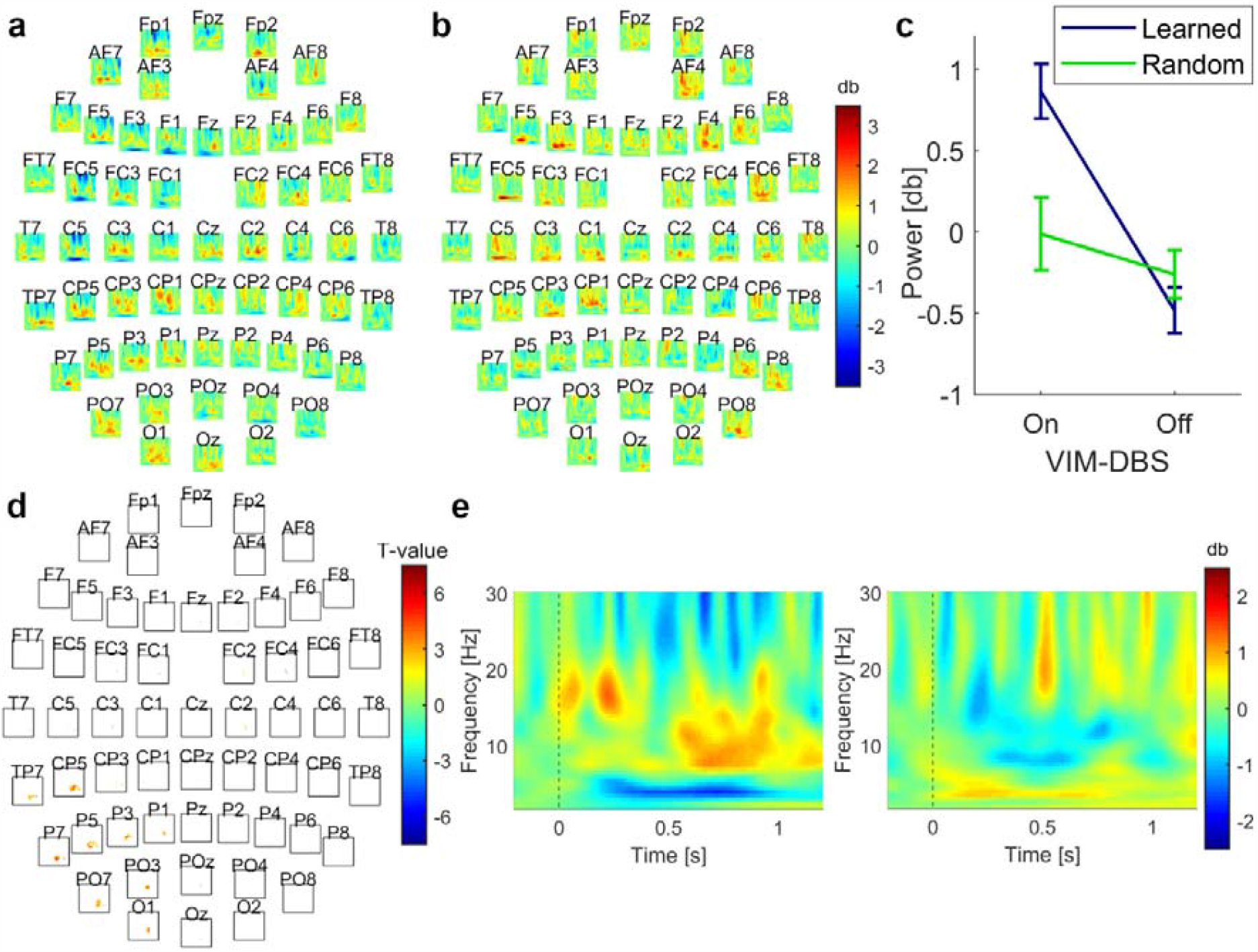
Changes in oscillatory power over the course of responding to a sequence (difference between Block 4 and Block 1). **a** Grand-average difference between DBS-ON and DBS-OFF for learned sequences. **b** Grand-average difference between DBS-ON and DBS-OFF for random trials. **c** Interaction for power values between *Stimulation Mode* and *Sequence Type*, showing mean and standard error. **d** Location of the observed cluster, representing significant difference between DBS-ON and DBS-OFF during learned sequences. The cluster included electrodes spanning the contralateral cortex over a central-parietal, parietal-occipital region. The channel contributing most to the cluster was CP5. **e** Left panel: channel CP5 during DBS-ON; right panel: channel CP5 during DBS-OFF. **a, b, d**: x-axis = Time: -200 to 1200 ms; y-axis = Frequency: 2-30 Hz.

These findings suggest a specific pattern of oscillatory power modulation in the alpha/beta bands when sequence learning occurred with DBS-ON, rather than a general performance improvement due to tremor reduction from VIM-DBS.

Across the course of learning, we observed an increase in alpha power in a cluster centered over left parietal cortex, contralateral to the moving hand. The contralateral parietal cortex contributes to motor sequence learning^15^ by integrating visual and somatosensory inputs^1,16^. Visuomotor control is established together with outputs to the dorsal PMC^16^, resulting in output to M1 for movement generation^1^. ET patients show decreased bidirectional functional connectivity between the parietal and contralateral motor cortex^7^. The increase in contralateral parietal alpha power observed when the stimulation was on may facilitate information integration between premotor and parietal areas as a sequence is learned through repetition.

The reduced alpha suppression we observed over central and ipsilateral motor cortical regions during VIM-DBS is consistent with the re-establishment of motor learning processes that were disrupted by pathological activity in this network. The effect was greater when the stimulation site was at more lateral aspects of the VIM, regions that are anatomically linked with cortical and subcortical motor structures^17^ and where single-cell recordings show tactile and kinesthetic responses^18^. Previous studies have shown that the alpha suppression over M1 typically observed during movement is attenuated when a movement sequence is repeated. This pattern has been hypothesized to reflect reduced attentional processing once the sequence was established^10,11^. On the other hand, a local increase in alpha band power has also been thought to have a time-dependent inhibitory effect on cortical functioning, specifically in regions no longer involved in task performance^19^. Future work will be needed to establish whether the reduction in alpha suppression during VIM-DBS indicates lower attentional requirements after a sequence is learned, or whether it reflects an enhancement of selective attention, with greater inhibition of task-relevant brain regions.

Beta band modulations may reflect cortical reorganization associated with sequence learning^10^. While beta suppression has been associated with diminishing interference effects by stabilizing the newly learned sequence during early consolidation^10^, prolonged beta band activity suppression is thought to hinder behavioral flexibility by promoting the maintenance of the current motor state, even if this is not optimal^20^. Greater beta suppression has been found in patients with Parkinson’s disease at the end of acquisition with diminished learning^12^. In our ET patient group, VIM-ON led to lower beta band suppression at the end of acquisition over the ipsilateral motor cortex and accompanied improved learning.

Interestingly, the reduction in alpha/beta sensorimotor suppression over motor cortex during stimulation was unilateral and ipsilateral to movement. Given that testing was limited to the right hand, we cannot say if this asymmetry (or the one noted in the parietal cortex) is related to the relationship of the cortex and moving hand (e.g., ipsilateral vs contralateral) or hemispheric specialization. It does suggest that the observed impact was on higher level processes associated with motor sequence learning rather than motor execution per se. In terms of hemispheric differences, previous work has shown the engagement of right PMC during spatial tasks as well as in the later stages of sequence learning^3^.

The effects of electrical stimulation on cognitive processing have been shown to depend on performing the relevant task during stimulation^21^. The current findings of a differential effect on oscillatory power, depending on whether task performance involved sequence learning, further supports this supposition.

The present results provide direct evidence for the engagement of the VIM in motor sequence learning, showing modulation of electrophysiological activity in nodes of the motor learning network through VIM-DBS^4^. We postulate that VIM-DBS interrupted pathophysiological activity in cortical networks, akin to the normalization of certain oscillatory patterns in M1 observed in Parkinsons’ disease patients who receive subthalamic nucleus stimulation DBS^22^. Subcortical, high frequency DBS appears to be able to override disease-related oscillatory patterns, promoting pro-kinetic oscillatory activity. Future work is required to establish whether DBS promotes re-establishment of normal oscillatory activity, suppresses abnormal activity, or imposes different activity patterns that enable improved function.

## 1. Materials and Methods

### 1.1. Participants

Sixteen right-handed patients with VIM electrodes previously implanted for DBS treatment of ET were recruited through the Stereotactic Neurosurgery Department, University Hospital, Magdeburg and were tested in an out-patient setting with EEG recorded while performing the experiment. Four datasets were excluded due to noisy EEG and lack of responses, resulting in the inclusion of N = 12 patients (Supplementary Material Table 2).

VIM-DBS electrode locations were determined relative to the anterior and posterior commissures (AC–PC line), based on co-registering the post-operative CT images and the electrode coordinates from the intraoperative stereotactic x-rays, with the pre-operative structural MRI images (Figure 1, Supplementary Material Table 2).

All participants gave written, informed consent prior to inclusion in the study, which was approved by the Local Ethics Committee of the Medical Faculty of the Otto von Guericke University Magdeburg, and carried out in accordance with the Declaration of Helsinki.

### 1.2. Study Design

Each participant performed two sessions of the SRTT on the same day, one session with (DBS-ON) and one without DBS (DBS-OFF), in a counterbalanced order. The task was presented using Presentation software (Neurobehavioral Systems, Berkeley, CA, USA).

Participants placed four fingers of their dominant (right) hand on four buttons of an ergonomically shaped response button pad. A row of four squares was shown on a computer screen. When a square turned red, participants were asked to respond by pressing the corresponding button on the response pad, with compatible S-R [stimulus–response] mapping, as quickly and accurately as possible. The squares were highlighted red according to a fixed, 12-element sequence at locations 1-3-2-1-4-1-2-3-1-3-2-4, or at random. The random presentation order was constrained, such that the same square was not highlighted consecutively, and each location was presented at least once every 12 items. The stimuli were presented for 500 ms and the inter-stimulus interval was fixed at 1200 ms, independent of the participants’ response times. Both sessions consisted of 4 blocks, each containing 144 trials. The 144 trials alternated between three cycles in which the stimuli followed a fixed sequence and three cycles in which the stimuli were selected at random, always starting with the sequential cycles.

### 1.3. DBS

Each DBS electrode probe had four contact locations at which stimulation could be applied. When the task was performed with the DBS on, stimulation was applied at the contact location, amplitude, frequency, and pulse width determined by the specialist DBS nurse to provide optimal tremor suppression while minimizing side-effects (Supplementary Material Table 2). This approach was taken for two reasons: the clinically-determined optimal parameters reflect the realistic impact on motor sequence learning in patients receiving VIM-DBS for tremor suppression, and from an ethical standpoint, only stimulation was applied that had been determined in routine clinical practice to provide maximal benefit to the patients. For statistical comparison, we estimated the total electrical energy delivered^25^ for both hemispheres.

Structural MRI data were recorded for clinical purposes using a Siemens Verio scanner (Siemens, Erlangen, Germany) equipped with a 32-channel head coil. The images were produced using the Inomed (Emmendingen, Germany) software package.

### 1.4. EEG Recording and Preprocessing

During the SRTT, EEG was recorded using a 64-electrode EEG cap (Brain Products, Gilching, Germany), with electrode AFz set as the ground and electrode FCz as the reference. The sampling rate was 500 Hz. EEG data were recorded with Brain Vision Recorder (Brain Products, Gilching, Germany) software, and the following offline data processing was conducted with MATLAB R2018a (MathWorks, Natick, MA, USA) and the toolboxes DBSFILT (version 0.18b)^14^ and Fieldtrip (version 20180826)^13^.

Raw data files were band-pass filtered from 1 Hz to 100 Hz and notch-filtered between 49 and 51 Hz using DBSFILT^14^. DBSFILT was then used to apply a Hampel filter to all datasets to remove aliasing peaks in lower frequencies, which are common DBS artifacts. Aliased frequencies can be detected following application of a Hampel filter, enabling removal of only the noise component at each interference frequency^14^. Further data processing was performed using Fieldtrip^13^. The data were segmented into epochs from -1400 ms before stimulus onset until 2600 ms after stimulus onset. The 200 ms before stimulus onset were used for baseline correction. Each highlighting of a square in red was considered a stimulus. Analysis of data time-locked to the stimulus is a common approach^5,12^ and offers the advantage that it resolves contamination by the activity related to the motor response, as well as tremor activity. The data were again band-pass filtered from 1 Hz to 30 Hz, using a padding length of 10 s. Bad channels were removed from the data based on visual inspection. Independent component analysis was performed to eliminate eye blinks and residual DBS artifacts, so that these trials could be retained. Afterwards, all epochs with values exceeding±100 _μ_V were excluded. Previously removed bad channels were then replaced using spherical spline interpolation. Surface Laplacians were calculated to reduce the effects of volume conduction, and to improve the spatial resolution of the EEG. All subsequent analyses were applied to the transformed data.

Time–frequency decomposition was performed through convolution with five-cycle complex Morlet wavelets for the frequencies ranging from 2 Hz to 30 Hz in increments of 0.5 Hz over the time window from -200 ms to 1200 ms, in increments of 10 ms. The time–frequency data were normalized to the baseline window 200 ms before until stimulus onset. Event-related spectral perturbations (ERSPs) were determined separately for each channel, block (Block 1, Block 4), stimulation mode (DBS-ON, DBS-OFF), sequence type (learned, random), and participant by averaging over the trials in which the target was correctly identified. To examine evolution of learning, a contrast was formed by subtracting participants’ individual ERSP in Block 1 from that in Block 4 separately for stimulation and sequence type.

For visualization, grand average ERSPs were then calculated across participants, and difference plots were created by subtracting the grand average DBS-OFF from DBS-ON.

### 1.5. Data Analysis and Statistics

#### 1.5.1. Behavioral analysis

Statistical analysis was performed using IBM SPSS Statistics 23 (IBM, Armonk, NY, USA). A repeated measures ANOVA with the within-subject factors *Stimulation Mode* (DBS-ON, DBS-OFF) and *Sequence Type* (Learned, Random) was applied to the reaction times, followed by post hoc two-sided paired T-tests.

#### 1.5.2. Cluster-based permutation tests

To identify effects of DBS on oscillatory spectral power, irrespective of task performance, and during the learned and random sequences, we performed three non-parametric cluster-based permutation tests^13^, including all channels, time points, and frequencies. The first was applied to all trials at the end of acquisition, including when the sequence was learned and when it was random. The second analyses were focused on the end of acquisition, where we assumed the sequence to be maximally learned in the condition with the repeated sequence,using all successfully learned trials during Block 4 to compare between DBS-ON and DBS-OFF conditions. To examine the time evolution of learning, we performed a third analysis, testing the contrast between Block 4 and Block 1 during the learned sequence and the random trials, and compared the DBS-ON against DBS-OFF conditions.

The CBPT enabled analysis of the neuronal data without a priori assumptions regarding the exact location or extent of a possible effect. The multiple comparison problem is resolved by applying a single test statistic to clusters (adjacent points in space, time, and frequency, which differ between conditions at a pre-defined threshold) instead of evaluating the differences between conditions at each sample point separately. Cluster formation was performed using a dependent samples two-sided t-test, with an uncorrected p-value threshold of p < 0.025 per side, and adjacency was defined as a minimum of two neighboring channels. A permutation distribution was obtained by pooling the averages per participant, irrespective of condition, then randomly assigning them to two categories using a Monte Carlo simulation. For each of 500 randomizations, t-tests were applied and clusters determined, with the sum of t-values of the maximum cluster per randomization as the cluster-based test statistic. The p-value was then derived by comparing the uncorrected observed cluster-based test statistic with the permutation distribution and was the proportion of randomizations in which the permuted cluster-based test statistic was larger than the observed cluster-based test statistic. P-values smaller than our critical alpha-level (p = 0.025 per side) were deemed significant. Effect size was calculated for the average of the cluster using Cohen’s 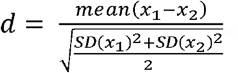.

#### 1.5.3 Post hoc testing

To confirm the specificity of the power differences between stimulation modes to sequence learning, the mean power values within the previously determined clusters were averaged over time, frequency, and electrodes during the repeated sequence, analogous power values were calculated from the data recorded during presentation of stimuli in a random order, and a post hoc repeated measures ANOVA with factors *Stimulation Mode* (DBS-ON, DBS-OFF) and *Sequence Type* (Learned, Random) was applied, followed by post hoc two-sided paired T-tests.

#### 1.5.4 Regression

A multiple linear regression was used to test whether clinical factors (right AC–PC coordinates, right total electrical energy delivered, and tremor score) predicted power at the end of acquisition. Power values during DBS-ON within the cluster were averaged over time, frequency, and electrodes.

## Notes

**Data availability statement:** The data that support the findings of this study are available from the corresponding authors upon reasonable request.

**Funding statement:** This work was supported by the Deutsche Forschungsgemeinschaft (DFG) [grant number SW 214/2-1 (CMSR)].

**Conflict of interest disclosure:** None.

### Competing Interest Statement

The authors have declared no competing interest.

### Summary of Updates

Further details relating to the statistics, including effect sizes and confidence intervals added.

