## Supplementary material for "Ventrointermediate thalamic stimulation improves motor learning in humans"

**Supplementary Table 1** Regression results using mean power with DBS-ON as the criterion

| | Unstandardized Coefficients | | Standardized Coefficients | | | 95% Confidence Interval | |
|---|---|---|---|---|---|---|---|
| Model | B | *SE* | Beta | *T* | Significance | LL | UL |
| Constant | 8.494 | 2.224 | | 3.820 | 0.009** | 3.053 | 13.935 |
| AC–PC-x (right) | -0.463 | 0.123 | -1.424 | -3.776 | 0.009** | -0.762 | -0.163 |
| AC–PC-y (right) | 0.259 | 0.099 | 0.681 | 2.615 | 0.040* | 0.017 | 0.501 |
| AC–PC-z (right) | 0.040 | 0.092 | 0.122 | 0.437 | 0.677 | -0.185 | 0.266 |
| TEED (right) | -0.026 | 0.008 | -0.775 | -3.303 | 0.047* | -0.045 | -0.588 |
| TRS-ON | 0.029 | 0.012 | 0.431 | 2.491 | 0.016* | 0.001 | 0.058 |

**Note:** [a] Dependent variable: mean power ($r$ = 0.941, $r^2$ = 0.886, Adj. $r^2$ = 0.791).
* indicates p < 0.05, ** indicates p < 0.01.

**Supplementary Table 2** Patient information and stimulation parameters. Values reported are the tremor rating score (TRS), while stimulation was on, and the stimulation type (m = monopolar; b = bipolar) for the left/right stimulating electrode. Stimulation frequency (Hz) was identical for both hemispheres. Amplitude (V = volts; mA = milliampere), pulse width (μs = microseconds), and electrode location are given separately for the left and right hemisphere. Electrode locations are given in relation to the AC–PC line (mm) lateral to the midline (x), posterior to the mid-commissural point (y) and inferior to the inter-commissural plane (z).

| | | | | | | Left hemisphere | | | | | Right hemisphere | | | | |
|---|---|---|---|---|---|---|---|---|---|---|---|---|---|---|---|
| | | | | | | | | AC–PC | | | | | AC–PC | | |
| ID | Gender | Age | TRS during DBS-ON | Stim Type | Frequency | Amplitude | Pulse width | x | y | z | Amplitude | Pulse width | x | y | z |
| 1 | f | 79 | 0 | m/m | 130 | 2.4 V | 60 | -11.5 | -7.0 | -3.2 | 2.0 V | 60 | 11.3 | -5.0 | -2.2 |
| 2 | m | 74 | 20 | m/m | 130 | 2 mA | 40 | -12.1 | -7.5 | -3.0 | 3.5 mA | 40 | 11.0 | -4.5 | -2.9 |
| 3 | f | 66 | 5 | b/b | 130 | 3.5 V | 60 | -13.1 | -2.5 | 1.4 | 3.0 V | 60 | 13.5 | -2.5 | 0.9 |
| 4 | m | 78 | 30 | m/m | 130 | 6.0 mA | 20 | -14.6 | -5.0 | 0.8 | 3.5 mA | 40 | 12.1 | -6.0 | -1.0 |
| 5 | m | 80 | 12 | m/m | 130 | 2.5 V | 60 | -12.5 | -4.0 | -2.5 | 2.5 V | 60 | 17.3 | 0.0 | 2.4 |
| 6 | f | 74 | 15 | m/m | 130 | 2.9 mA | 60 | -15.3 | -5.0 | 0.6 | 3.0 mA | 60 | 14.6 | -3.0 | 1.8 |
| 7 | m | 73 | 9 | m/m | 130 | 3.25 mA | 40 | -12.2 | -5.5 | -2.1 | 3.0 mA | 40 | 14.9 | -4.5 | -0.5 |
| 8 | m | 77 | 13 | b/b | 130 | 2.0 V | 60 | -13.2 | -4.5 | 0.1 | 2.8 V | 60 | 13.3 | -5.5 | 1.7 |
| 9 | f | 60 | 0 | m/b | 130 | 2.0 V | 90 | -12.1 | -4.0 | -0.8 | 2.5 V | 90 | 10.5 | -4.5 | -1.0 |
| 10 | m | 78 | 20 | m/m | 130 | 1.8 V | 60 | -14.7 | -4.5 | 1.4 | 2.6 V | 60 | 12.9 | -5.5 | 1.5 |
| 11 | f | 62 | 1 | m/m | 130 | 2.8 V | 60 | -12.4 | -3.5 | 0.1 | 3.6 V | 60 | 11.8 | -3.5 | 1.3 |
| 12 | f | 67 | 6 | m/m | 130 | 2.6 mA | 40 | -12.4 | -2.5 | -1.5 | 2.6 mA | 40 | 12.2 | -4.5 | -2.8 |
